# Genetic profiling of Vietnamese population from large-scale genomic analysis of non-invasive prenatal testing data

**DOI:** 10.1101/868588

**Authors:** Ngoc Hieu Tran, Thanh Binh Vo, Van Thong Nguyen, Nhat Thang Tran, Thu-Huong Nhat Trinh, Hong-Anh Thi Pham, Thi Hong Thuy Dao, Ngoc Mai Nguyen, Yen-Linh Thi Van, Vu Uyen Tran, Hoang Giang Vu, Quynh-Tram Nguyen Bui, Phuong-Anh Ngoc Vo, Huu Nguyen Nguyen, Quynh-Tho Thi Nguyen, Thanh-Thuy Thi Do, Phuong Cao Thi Ngoc, Dinh Kiet Truong, Hoai-Nghia Nguyen, Hoa Giang, Minh-Duy Phan

## Abstract

The under-representation of several ethnic groups in existing genetic databases and studies have undermined our understanding of the genetic variations and associated traits or diseases in many populations. Cost and technology limitations remain the challenges in performing large-scale genome sequencing projects in many developing countries, including Vietnam. As one of the most rapidly adopted genetic tests, non-invasive prenatal testing (NIPT) data offers an alternative untapped resource for genetic studies. Here we performed a large-scale genomic analysis of 2,683 pregnant Vietnamese women using their NIPT data and identified a comprehensive set of 8,054,515 single-nucleotide polymorphisms, among which 8.2% were new to the Vietnamese population. Our study also revealed 24,487 disease-associated genetic variants and their allele frequency distribution, especially 5 pathogenic variants for prevalent genetic disorders in Vietnam. We also observed major discrepancies in the allele frequency distribution of disease-associated genetic variants between the Vietnamese and other populations, thus highlighting a need for genome-wide association studies dedicated to the Vietnamese population. The resulted database of Vietnamese genetic variants, their allele frequency distribution, and their associated diseases presents a valuable resource for future genetic studies.

## Background

Following the successful initiative of the 1000 genomes project [1], several large-scale genome and exome sequencing projects have been conducted, either as international collaboration efforts such as ExACT [2], gnomAD [3], or for a specific country or population [4–8]. Those projects have provided comprehensive profiles of human genetic variation in some populations, paving the way for unprecedented advance in treatment of common genetic diseases. However, the lack of diversity and the under-representation of several populations in genome sequencing projects and genome-wide association studies (GWAS) have increasingly become a critical problem [9,10]. For instance, Gurdasani *et al.* found that the representation of ethnic groups in GWAS was significantly biased, with nearly 78% of the participants having European ancestries, whereas the two major populations, Asian and African, only accounted for 11% and 2.4%, respectively [10]. Vietnam has a population of 96.5 million, the 15^th^ highest in the world and the 9^th^ highest in Asia. Yet there was merely one dataset of 99 Vietnamese individuals that had been studied as part of the 1000 genomes project (population code KHV, the Kinh ethnic group in Ho Chi Minh City, Vietnam). A recent study has sequenced genomes and exomes of another 305 individuals to further expand the Vietnamese genetic database [11]. However, costs and technologies to perform large-scale genome sequencing projects still remain a challenge for most developing countries, including Vietnam.

An alternative approach has been proposed recently to re-use the low-coverage genome sequencing data from non-invasive prenatal testing (NIPT) for large-scale population genetics studies [12,13]. NIPT is a method that sequences cell-free DNA from maternal plasma at an ultra-low depth of 0.1-0.2x to detect fetal aneuploidy [14]. By combining a sufficiently large number of NIPT samples, one could obtain a good representation of the population genetic variation. The benefits of re-using NIPT data for population genetics are manifold. First, the data can be re-used at no extra cost given the approval and consent of the participants. As one of the most rapidly adopted genetic tests, NIPT has been successfully established and become a standard screening procedure with thousands to millions of tests performed world-wide, including many developing countries such as Vietnam [14]. Using NIPT data for population genetic studies may also reduce privacy concerns since the genetic variants can only be analyzed by aggregating a large number of samples and the results can only be interpreted at the population level. A single sample tells little about the genetic information of an individual due to low sequencing depth. Last but not least, previous studies have suggested that sequencing a large number of individuals at a low depth might provide more accurate inferences of the population genetic structure than the traditional approach of sequencing a limited number of individuals at a higher depth, especially when the budget is limited [15,16].

In this paper, we presented the first study of Vietnamese genetic variations from non-invasive prenatal testing data, and to the best of our knowledge, the third of such kinds in the world [12,13]. We analyzed the NIPT data of 2,683 pregnant Vietnamese women to identify genetic variants and their allele frequency distribution in the Vietnamese population. We also studied the relationships between the Vietnamese genetic profile and common genetic disorders, and discovered pathogenic variants related to prevalent diseases in Vietnam. Finally, we highlighted the differences in the distribution of disease-associated genetic variants between the Vietnamese and other populations, thus highlighting a need for genome-wide association studies dedicated to the Vietnamese population. The resulted database of Vietnamese genetic variants from NIPT data is made available to facilitate future research studies in population genetics and associated traits or diseases.

## Results

### Data collection

A total of 2,683 pregnant Vietnamese women who performed non-invasive prenatal testing during the period from 2018-2019 at the Medical Genetics Institute, Vietnam, were recruited to the study. The participants have approved and given written consent to the anonymous re-use of their genomic data for the study. All information of the participants is confidential and not available to the authors, except the records that their NIPT and pregnancy results are normal. The study was approved by the institutional ethics committee of the University of Medicine and Pharmacy, Ho Chi Minh city, Vietnam. The whole genome of each participant was sequenced to an average of 3.6 million paired-end reads of 2×75 bp, which corresponds to a sequencing depth of 0.17x per sample.

### Genome coverage and sequencing depth of the NIPT dataset

Data pre-processing was first performed on each of 2,683 samples and the results were stored in binary alignment map (BAM) format, one BAM file per sample. The data pre-processing steps include: quality control of raw data using FastQC [17], trimmomatic [18]; alignment of paired-end reads to the human reference genome (build GRCh38) using bwa [19], samtools [20], MarkDuplicates [21]; and summary of alignment results using Qualimap [22], bedtools [23], IGV [24]. The quality of raw data and alignment results are presented in Supplementary Figures S1 and S2. Raw data showed high sequencing quality, no error or bias was observed. The mapping quality and insert size distributions followed closely what expected across the reference genome. The overall sequencing error rate was about 0.3%. More details of the data pre-processing steps can be found in the Methods section.

The average genome coverage and depth were 14.59% and 0.17x per sample, respectively, and aggregated to 95.09% and 462x across 2,683 samples (Figure 1a). Although the sequencing depth per sample was low, there might be more than one read from the same sample overlapping at a genome position. This problem may affect the estimation of allele frequency because the estimation is based on the assumption that a sample may contribute only 0 or 1 allele (read) at any given genome position [12,13]. For instance, we found that the average percentage of genome positions with depth 2x (i.e. covered by two overlapping reads) in a sample was 1.75% (Figure 1a). These overlapping reads occurred randomly across the reference genome and the samples. At any genome position, there were on average 47 out of 2,683 samples that each contributed two reads (Supplementary Figure S3). To address this problem, we followed a filtering strategy from previous studies [12,13] to keep only one read if there were overlapping reads in a sample. Thus, for every genome position, each sample could only contribute up to one read, and when the samples were aggregated, all reads at any position were obtained from different samples. In addition, we also removed alignments with low mapping quality scores (MAPQ < 30).

**Figure 1.**
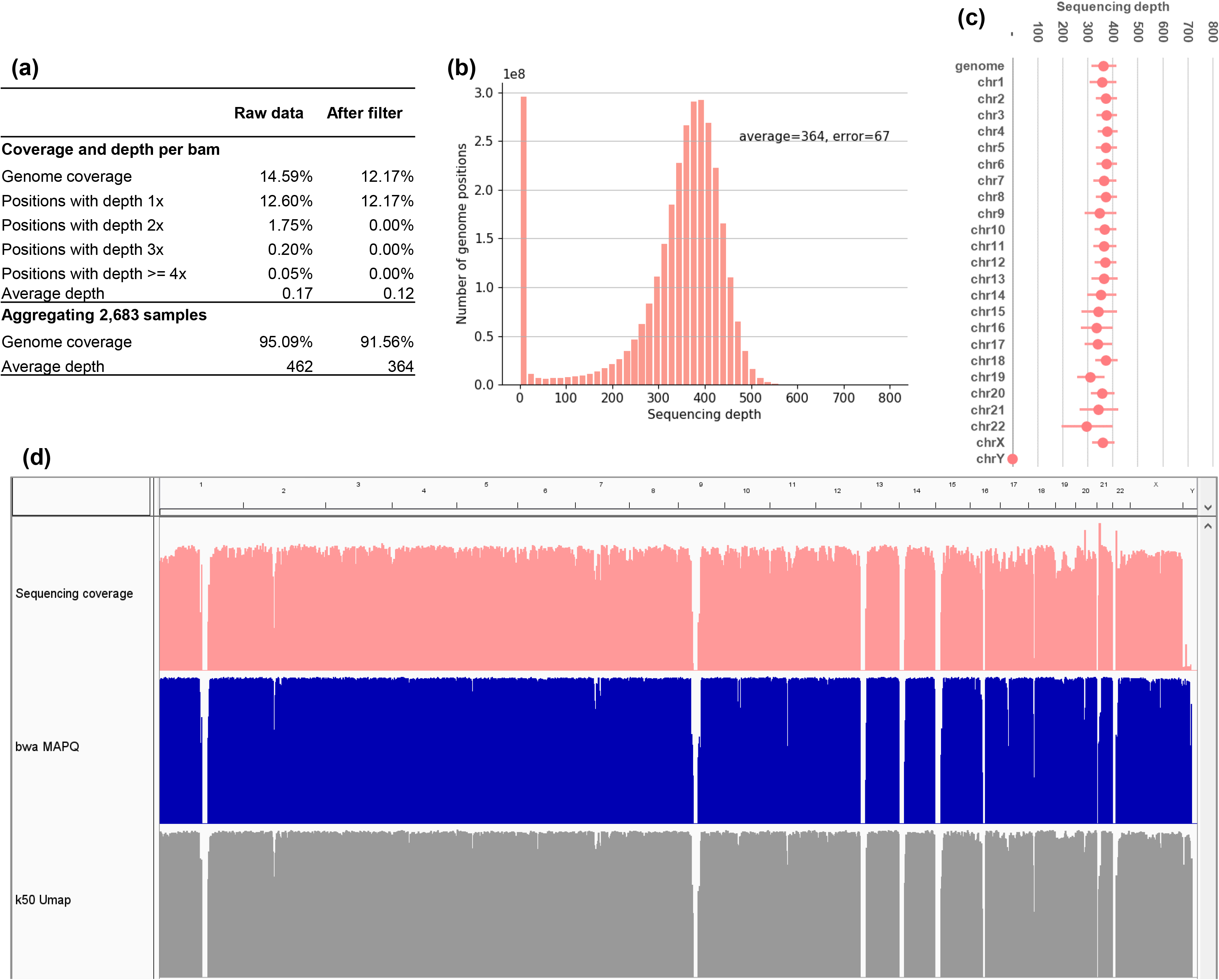
Distributions of genome coverage and sequencing depth of the NIPT dataset. (a) Average genome coverage and sequencing depth per sample and from all samples combined. (b) Summary histogram of sequencing depth over all genome positions. (c) Distribution of sequencing depth per chromosome. (d) IGV tracks of sequencing depth, bwa MAPQ score, and Umap k50 mappability across the whole genome.

After the filtering step, the sequencing depth was reduced from 0.17x to 0.12x per sample. The aggregated sequencing depth of 2,683 samples was 364x and the genome coverage was 91.56% (Figure 1a). The distributions of genome coverage and sequencing depth are presented in Figures 1b-d. The sequencing depth was approximately uniform across the reference genome, except for low-mappability regions and chromosome Y. The distributions of sequencing depth and MAPQ score also closely followed the mappability of the human reference genome obtained from Umap [25]. The average sequencing depth of chromosome Y was about 2.1% that of the whole genome, consistent with the proportion of fetal DNA in NIPT samples (8-10%) [14].

### Variant calling and validation

We aggregated 2,683 samples into one and used Mutect2 from GATK [26,27] for variant calling and allele frequency estimation. In addition to its main function of somatic calling, Mutect2 can also be used on data that represents a pool of individuals, such as our NIPT dataset, to call multiple variants at a genome site [12,13,28]. The called variants were further checked against strand bias, weak evidence, or contamination using FilterMutectCalls. The allele frequencies were estimated based on the numbers of reads aligned to the reference and the alternate alleles. For validation, we compared our NIPT call set to the KHV (Kinh in Ho Chi Minh City, Vietnam) and EAS (East Asian) populations from the 1000 genomes project [1], as well as the dbSNP database (version 151, [29]).

We identified a total of 8,054,515 SNPs from the NIPT dataset. The transition to transversion ratio was 2.0 over the whole genome and 2.8 over protein coding regions, which was similar to the observed ratios from previous genome or exome sequencing projects. As expected, a majority of these SNPs, 7,390,020 or 91.8%, had been reported earlier in the KHV call set (Figure 2a). Since the KHV population only had 99 individuals, we further looked into its common SNPs that were shared by at least two individuals. We found that the NIPT call set recovered 90.5% of the KHV common SNPs (Supplementary Figure S4; 6,889,016 / 7,609,526 = 90.5%). This sensitivity is in line with the genome coverage reported earlier in Figure 1a. An important advantage of NIPT data is the ability of sampling a large number of individuals to better represent a population and to accurately estimate the allele frequency. We found a strong Pearson correlation of 98.8% between the allele frequency of the NIPT call set and that of the KHV call set (Figure 2b). Furthermore, thanks to its larger sample size, the NIPT allele frequency indeed showed better resolution than the KHV one, as evidenced by vertical trails in Figure 2b or a zoomed-in view in Supplementary Figure S5.

**Figure 2.**
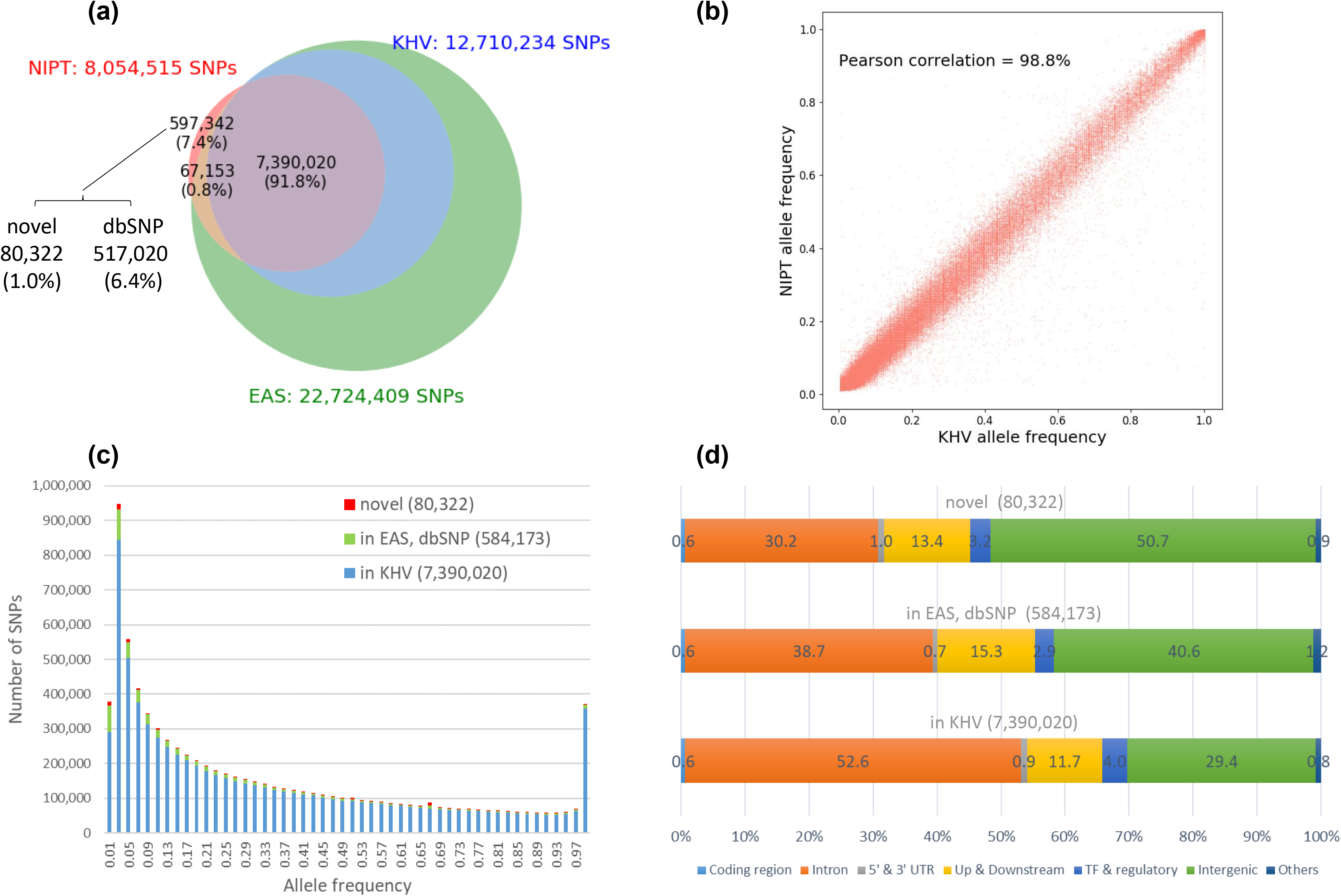
Summary of the NIPT call set. (a) Venn diagram comparison between the NIPT call set, the KHV and EAS call sets from the 1000 genomes project, and the dbSNP database. The percentages were calculated with respect to the NIPT call set. (b) Scatter plot comparison of allele frequency estimated from the NIPT and the KHV call sets. (c) Allele frequency distribution of the NIPT call set. (d) Distribution of locations and effects of variants in the NIPT call set.

Our NIPT call set included 664,495 (8.2%) SNPs that had not been reported in the KHV call set. Among them, 67,153 (0.8%) were found in the EAS call set, another 517,020 (6.4%) were found in the dbSNP database, and the remaining 80,322 (1.0%) were novel SNPs (Figure 2a). Majority of those SNPs had allele frequencies less than 10%. The overall allele frequency distribution of our NIPT call set is presented in Figure 2c.

We used VEP (Variant Effect Predictor [30]) to analyze the effects of 8,054,515 variants in the NIPT call set (Figure 2d). About 1.5% of the SNPs were located in the coding and UTR regions, 12% in the upstream and downstream regions, and 4% in the TF regulatory regions. More than 80% of the SNPs were located in the intron and intergenic regions. We also noted that the new SNPs and those in KHV had similar proportions of coding, UTR, upstream and downstream, and TF regulatory regions (Figure 2d).

### Analysis of pathogenic SNPs and their allele frequencies in Vietnamese population

We searched the NIPT call set against the ClinVar database (version 20191105, [31]) to explore the associations between Vietnamese genomic variants and common genetic diseases. We identified 24,487 SNPs with ClinVar annotations that have been reviewed by at least one research group (Table 1). Among them, five SNPs were classified as “Pathogenic” or “Likely pathogenic”, 117 SNPs were found to affect a “drug response”, and a majority of the remaining were classified as “Benign” or “Likely benign”. We also noted that 391 ClinVar-annotated SNPs (1.6%), including 1 pathogenic SNP, had not been reported in the KHV call set.

**Table 1.** Summary of ClinVar annotations for the NIPT call set.

| ClinVar clinical significance | in KHV | not in KHV |
| --- | --- | --- |
| Benign or Likely benign | 23,414 | 300 |
| Uncertain significance | 353 | 74 |
| Pathogenic or Likely pathogenic | 4 | 1 |
| Drug response | 114 | 3 |
| Others | 211 | 13 |
| **Total number of annotations** | **24,096** | **391** |

Table 2 and Supplementary Table S1 present the details of five pathogenic SNPs identified in our NIPT call set. Their associated genetic diseases include: erythropoietic protoporphyria, non-syndromic genetic deafness, Joubert syndrome, hemochromatosis type 1, and 5-alpha reductase deficiency. The SNP rs9332964 C>T in *SRD5A2*, which is associated with 5-alpha reductase deficiency, had not been reported in the KHV call set. 5-alpha reductase deficiency is an autosomal recessive disorder that affects male sexual development. This SNP is rare in the world and East Asia populations, but was found to be more common in the Vietnamese population (allele frequency of 0.05%, 0.67%, and 2.90%, respectively). This SNP had also been reported in a recent study [11] at a very low allele frequency of 1.36%.

**Table 2.** Pathogenic variants identified from the NIPT call set.

| Variant information |  |  |  |  | ClinVar annotations |  |  | Allele frequency |  |  |
| --- | --- | --- | --- | --- | --- | --- | --- | --- | --- | --- |
| chr | position | dbSNP | Ref | Alt | ID | Gene | Conditions | NIPT | gnomAD EAS | gnomAD |
| chr18 | 57571588 | rs2272783 | A | G | 562 | FECH | Erythropoietic protoporphyria | 28.10% | 32.57% | 11.23% |
| chr13 | 20189473 | rs72474224 | C | T | 17023 | GJB2 | Nonsyndromic hearing loss and deafness | 13.40% | 8.35% | 0.76% |
| chr13 | 72835359 | rs17089782 | G | A | 217689 | PIBF1 | Joubert syndrome | 6.80% | 5.13% | 1.36% |
| chr6 | 26090951 | rs1799945 | C | G | 10 | HFE | Hemochromatosis type 1 | 5.10% | 3.41% | 10.82% |
| chr2 | 31529325 | rs9332964 | C | T | 3351 | SRD5A2 | 5-alpha reductase deficiency | 2.90% | 0.67% | 0.05% |

We noticed that the allele frequencies of the five pathogenic SNPs varied considerably between the Vietnamese, the East Asia, and the world populations (Table 2). For instance, the SNP rs72474224 C>T in *GJB2* is commonly liked to non-syndromic hearing loss and deafness, which is also the most prevalent genetic disorder in the Vietnamese population. We found that its allele frequency in the Vietnamese population was 60% higher than in the East Asia population, which in turn was an order of magnitude higher than in the world population (13.40%, 8.35%, and 0.76%, respectively). The allele frequency was consistent with the estimated carrier frequency of 1 in 5 in the Vietnamese population. Similarly, the allele frequency of rs2272783 A>G in *FECH*, which is associated with erythropoietic protoporphyria, was nearly three times higher in the Vietnamese and East Asia populations than in the world population (28.10%, 32.57%, and 11.23%, respectively). The prevalence of this pair of SNP and disease in East and Southeast Asia has been reported previously in [32]. On the other hand, the allele frequency of rs1799945 C>G in *HFE*, which is associated with hemochromatosis type 1, was about two and three times lower in the Vietnamese and East Asia populations than in the world population (5.10%, 3.41%, and 10.82%, respectively). Such discrepancies were also observed for “Benign” variants, e.g., those related to autosomal recessive non-syndromic hearing loss (Supplementary Table S2). The variations strongly suggest that population-specific genome-wide association studies are required to provide a more accurate understanding of the clinical significance of genetic variants and the true disease prevalence in the Vietnamese population.

## Discussion

In this study, we analyzed the genomes of 2,683 pregnant Vietnamese women from their non-invasive prenatal testing data. The genomes were originally sequenced at a low depth of approximately 0.17x per sample for the purpose of fetal aneuploidy testing [14]. Here we combined the 2,683 samples to a total sequencing depth of 364x and performed variant calling and analysis for the Vietnamese population. We identified a comprehensive set of 8,054,515 SNPs at a high level of sensitivity and accuracy: 90.5% of Vietnamese common SNPs were recovered; 99% of identified SNPs were confirmed in existing databases; and a strong correlation of 98.8% to known allele frequencies. The results were exciting given that the total sequencing depth of our dataset, 364x, was merely equivalent to sequencing 20 individuals at a moderate depth of 20x. It also suggests that there is still plenty of room for improvement by increasing the number of NIPT samples. For instance, Liu *et al.* have demonstrated a large-scale population genetic analysis based on hundreds of thousands of NIPT samples for the Chinese population [12].

Another benefit of using NIPT data is cost-effective. In our study, the dataset was re-used at no cost with written consent from the participants. The whole analysis pipeline was done within a week on a cloud computing platform for a few hundred dollars. Thus, the overall cost was negligible compared to that of a typical genome sequencing project. The cost advantage of this approach may play a major role in large-scale genome sequencing projects, especially in developing countries where technologies and resources are still limited.

Our study revealed 24,487 disease-associated genetic variants, especially five pathogenic variants for prevalent genetic disorders in the Vietnamese population. We also found major discrepancies in the allele frequency distribution of genetic variants between the Vietnamese, the East Asia, and the world populations. Thus, a comprehensive genetic profile and genome-wide association studies dedicated to the Vietnamese population are highly desired. Knowing the distribution of genetic disorders in the population will be useful for public health policy and planning, preventive medicine, early genetic screening strategies, etc.

Some technical and design limitations in our study could be addressed in future research to improve the application of NIPT data in population genetics studies. First, currently there is no variant calling tool that is designed specifically for NIPT data. Here we used Mutect2 and previous studies also used similar somatic calling tools with the purpose of identifying all possible variants at a genome site [12,13]. Thus, we took a conservative approach to consider only SNPs but not indels to ensure a reliable call set. Another limitation was the exclusion of chromosome Y due to its low coverage as a result of limited amount of fetal DNA in NIPT samples. This problem could be addressed by increasing the number of samples to obtain enough sequencing coverage for reliable variant calling. NIPT data is also biased by sex, with only ~5% of the data coming from male population (assuming a 10% cell-free fetal DNA fraction with 50% male fetuses). In general, sufficiently large sample size is essential to use NIPT data for population genetics research. Thus, privacy policy, code of ethics, and standards of practice need to be established to protect the confidential information and data of the participants.

## Conclusions

We showed that non-invasive prenatal testing data could be reliably used to reconstruct the genetic profile of the Vietnamese population. Our study identified pathogenic variants for prevalent genetic diseases in the Vietnamese population and called for a need for population-specific genome-wide association studies. The resulted database presents a valuable resource for future studies of genetic variations and associated traits or diseases, not only for the Vietnamese population but also for other Southeast-Asia and Asia populations. Our results also demonstrated that non-invasive prenatal testing data provides a valuable and cost-effective resource for large-scale population genetic studies.

## Methods

### Sample preparation

Cell-free DNA (cfDNA) in maternal plasma was extracted using MagMAX Cell-Free DNA Isolation Kit from Thermo Fisher Scientific (Waltham, MA, USA). Library preparation was done using NEBNext Ultra II DNA Library Prep Kit from New England BioLabs (Ipswich, MA, USA). The samples were sequenced on the NextSeq 550 platform using paired-end 2×75 bp Reagent Kit from Illumina (San Diego, CA, USA).

### Bioinformatics analysis pipeline

Quality check of raw sequencing data was performed using FastQC (version 0.11.8, [17]). Paired-end reads were trimmed to 75 bp, adapters (“TruSeq3-PE-2.fa”) and low-quality bases were removed using trimmomatic (version 0.39, [18]). We only kept pairs with both reads surviving the trimming (about 95.7% of the dataset). The reads were then aligned to the human reference genome, build GRCh38 (hg38), using bwa mem (version 0.7.17-r1188, [19]). Supplementary hits were marked as secondary for Picard compatibility. Alignment results were sorted and indexed using samtools (version 1.9, [20]). Potential PCR duplicates were marked using MarkDuplicates from GATK (version 4.1.1.0, [26]).

Alignments with mapping quality scores less than 30 were discarded. In-house Python scripts were developed to mark overlapping alignments and to keep only one of them. Qualimap (version 2.2.1, [22]), bedtools (version 2.25.0 [23]), and IGV (version 2.4.19, [24]) were used to summarize the alignment results and to calculate genome coverage and sequencing depth.

Mutect2 from GATK (version 4.1.1.0 [26,27]) was used in tumor-only mode for variant calling. All samples were assigned the same sample name to combine them before variant calling. FilterMutectCalls was used to exclude variants with weak evidence, strand bias, or contamination. bcftools (version 1.9, [33]) was used to filter, summarize, and compare VCF (Variant Call Format) files. VEP (version 98, [30]) was used to predict the effects of variants and to annotate them against dbSNP (version 151, [29]) and ClinVar (version 20191105, [31]) databases.

The whole analysis pipeline and parameter settings can be found in the attached Python scripts.

## Declarations

### Ethics approval and consent to participate

The study was approved by the institutional ethics committee of the University of Medicine and Pharmacy, Ho Chi Minh city, Vietnam. The participants who performed NIPT triSure at Medical Genetics Institute, Vietnam, have approved and given written consent to the anonymous re-use of their genomic data for this study.

### Consent for publication

All authors have read and approved the manuscript for publication.

### Availability of data and materials

The database of genetic variants identified from NIPT data and the Python scripts for bioinformatics analysis pipeline is available on GitHub: https://github.com/nh2tran/NIPT_WGS

### Authors’ contributions

TBV, HATP, HTD, NMN, YLTV, VUT, HGV, QTNB, PANV performed experiments.

VTN, NTTM, THNT, TTTD recruited patients and performed clinical analysis.

HNN, QTTN, PCTN, DKT designed experiments and analyzed data.

HNN supervised the project.

NHT, HG, MDP analyzed the data and wrote the manuscript, designed experiments and analyzed sequencing data.

## Acknowledgements

The authors thank Dr. Kwok Pui Choi for critical reading of the manuscript.

