## Supplementary Materials for "Genetic profiling of Vietnamese population from large-scale genomic analysis of non-invasive prenatal testing data"

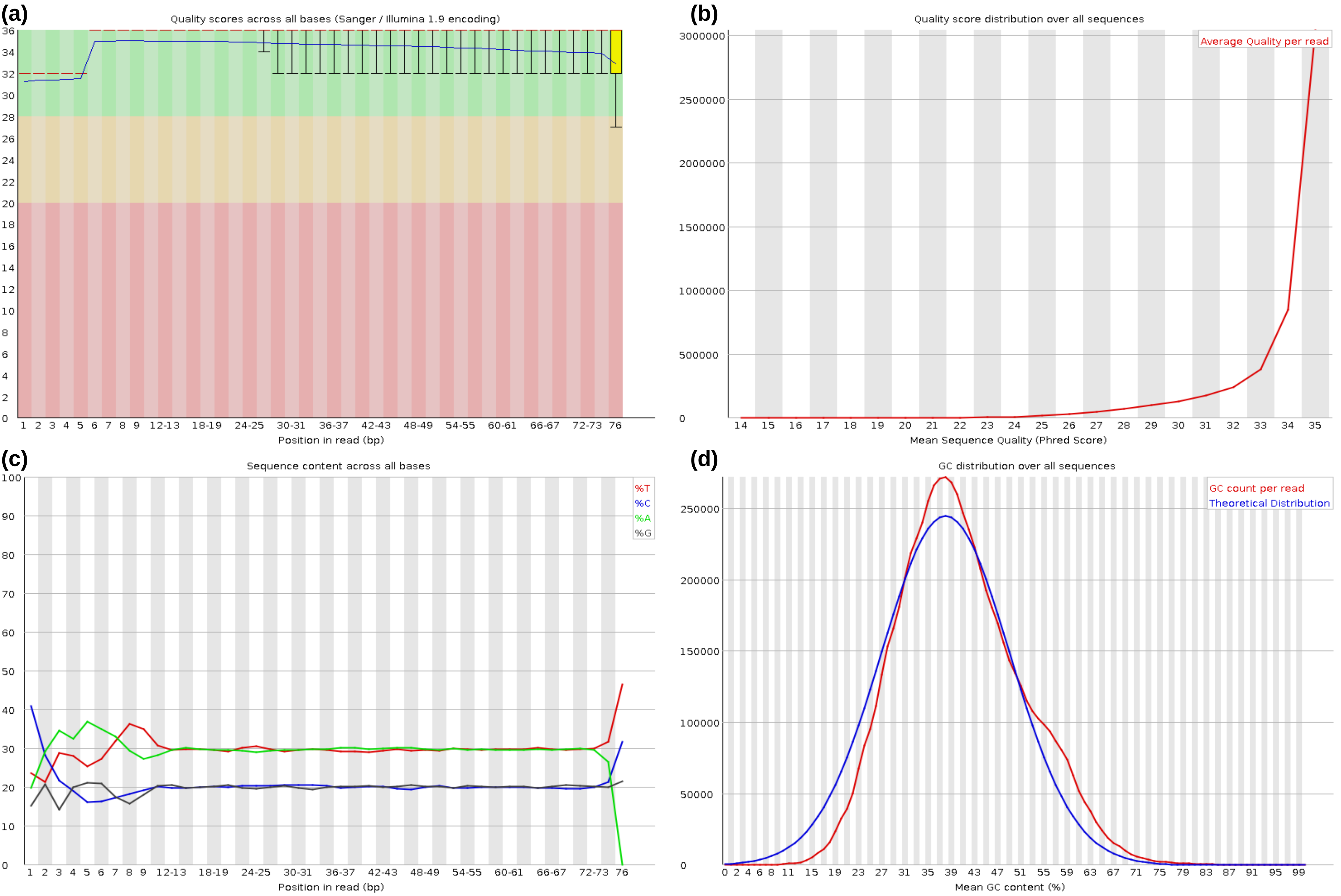


Figure S1. Quality control check on raw sequence data by FastQC. (a) Per base sequence quality. (b) Per sequence quality scores. (c) Per base sequence content. (d) Per sequence GC content.

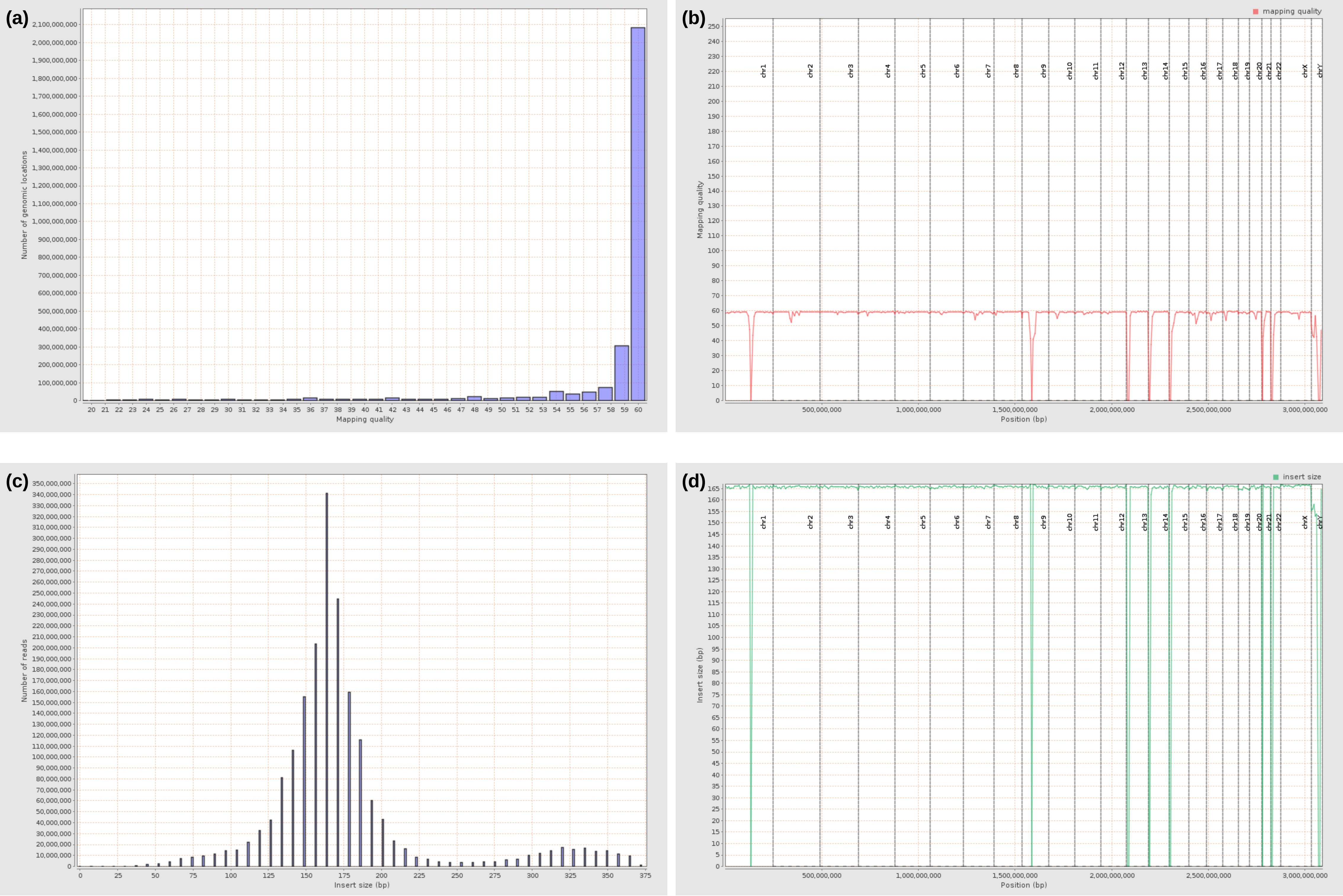


Figure S2. Qualimap report for the quality control of alignment results. (a) Mapping quality histogram. (b) Mapping quality across reference. (c) Insert size histogram. (d) Insert size across reference.

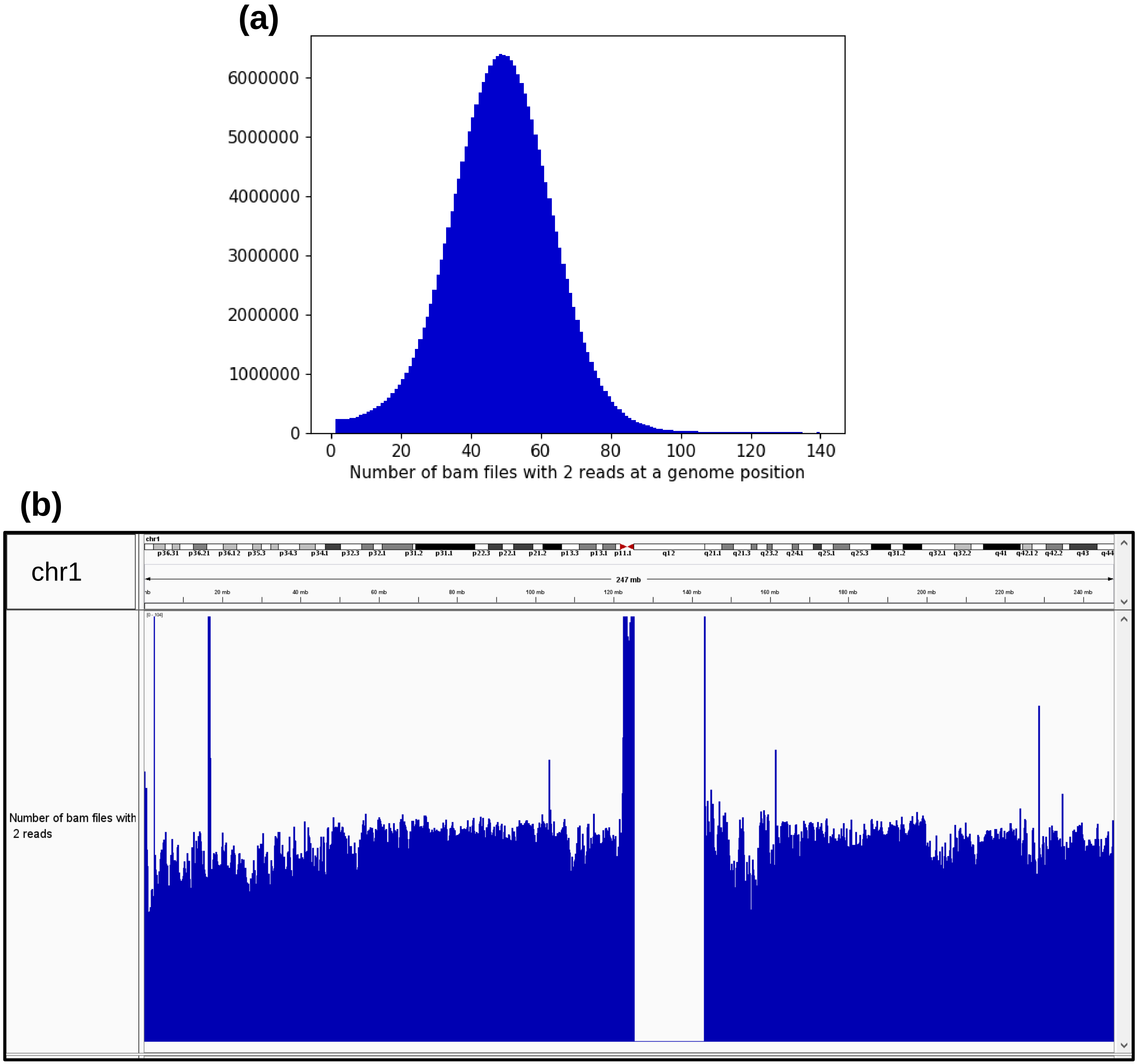


Figure S3. Distribution of the number of bam files that each contributed 2 reads at a genome position. (a) Summary histogram over all genome positions (y-axis shows the number of positions). (b) Distribution along chromosome 1.

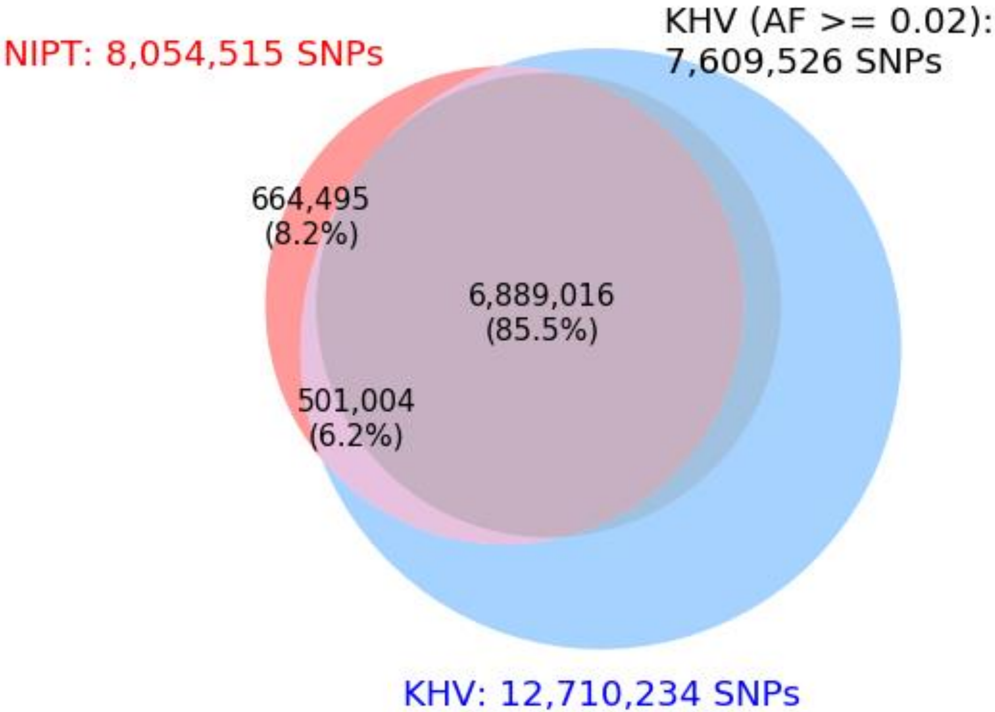


Figure S4. Venn diagram comparison between the NIPT call set, the KHV call set and its subset with allele frequency of at least 2%. The percentages were calculated with respect to the NIPT call set.

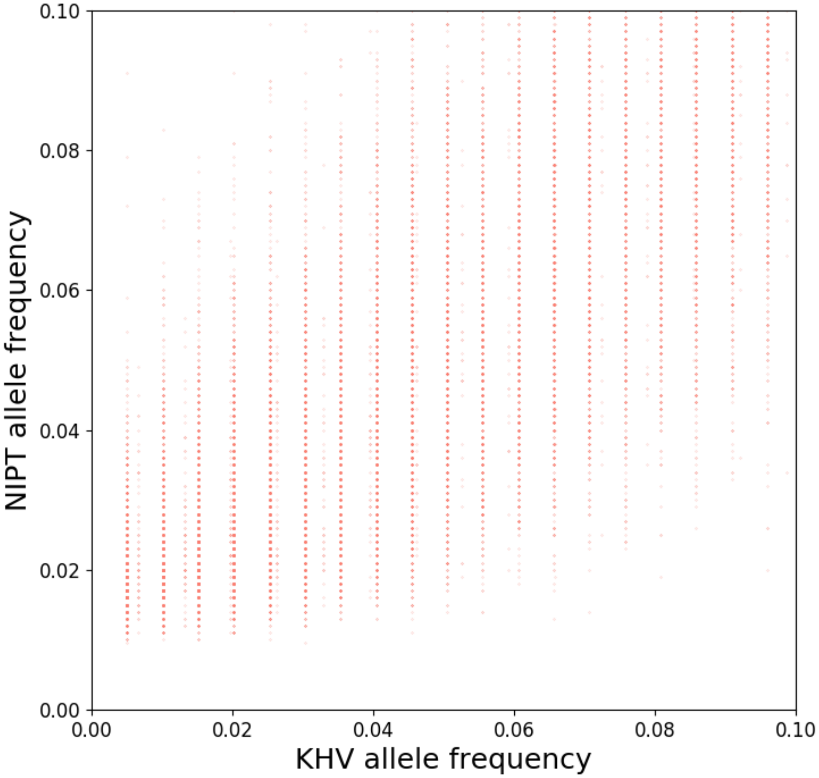


Figure S5. NIPT estimation of allele frequency showed better resolution than KHV's thanks to larger sample size.

**Table S1. Pathogenic variants identified from the NIPT call set and their allele frequencies in the KHV population**

| Variant information | | | | | ClinVar annotations | | | Allele frequency | | |
|---|---|---|---|---|---|---|---|---|---|---|
| chr | position | dbSNP | Ref | Alt | ID | Gene | Conditions | NIPT | KHV | Le *et al.* [11] |
| chr18 | 57571588 | rs2272783 | A | G | 562 | FECH | Erythropoietic protoporphyria | 28.10% | 25.76% | 22.20% |
| chr13 | 20189473 | rs72474224 | C | T | 17023 | GJB2 | Nonsyndromic hearing loss and deafness | 13.40% | 8.59% | 9.64% |
| chr13 | 72835359 | rs17089782 | G | A | 217689 | PIBF1 | Joubert syndrome | 6.80% | 4.55% | 6.33% |
| chr6 | 26090951 | rs1799945 | C | G | 10 | HFE | Hemochromatosis type 1 | 5.10% | 4.55% | 3.82% |
| chr2 | 31529325 | rs9332964 | C | T | 3351 | SRD5A2 | 5-alpha reductase deficiency | 2.90% | not found | 1.36% |

**Table S2. ClinVar annotations of SNPs identified on gene GJB2.**

| Variant information | | | | | ClinVar annotations | | | | Allele frequency | | |
|---|---|---|---|---|---|---|---|---|---|---|---|
| chr | position | dbSNP | Ref | Alt | ID | Gene | Conditions | Interpretation | NIPT | gnomAD EAS | gnomAD |
| chr13 | 20188817 | rs3751385 | A | G | 36277 | GJB2 | Nonsyndromic hearing loss, recessive | Benign | 73.20% | 55.56% | 74.86% |
| chr13 | 20189503 | rs2274084 | C | T | 36279 | GJB2 | Nonsyndromic hearing loss, recessive | Benign | 15.50% | 27.80% | 5.04% |
| chr13 | 20189473 | rs72474224 | C | T | 17023 | GJB2 | Nonsyndromic hearing loss, recessive | Pathogenic | 13.40% | 8.35% | 0.76% |
| chr13 | 20189241 | rs2274083 | T | C | 44739 | GJB2 | Nonsyndromic hearing loss, recessive | Benign | 8.90% | 18.81% | 1.46% |
| chr13 | 20188974 | rs76838169 | A | G | 44762 | GJB2 | Nonsyndromic hearing loss, recessive | Benign | 3.30% | 5.91% | 0.42% |
